# *De novo* computational RNA modeling into cryoEM maps of large ribonucleoprotein complexes

**DOI:** 10.1101/332791

**Authors:** Kalli Kappel, Shiheng Liu, Kevin P. Larsen, Georgios Skiniotis, Elisabetta Viani Puglisi, Joseph D. Puglisi, Z. Hong Zhou, Rui Zhao, Rhiju Das

## Abstract

RNA-protein assemblies carry out many critical biological functions including translation, RNA splicing, and telomere extension. Increasingly, cryo-electron microscopy (cryoEM) is used to determine the structures of these complexes, but nearly all maps determined with this method have regions in which the local resolution does not permit manual coordinate tracing. Because RNA coordinates typically cannot be determined by docking crystal structures of separate components and existing structure prediction algorithms cannot yet model RNA-protein complexes, RNA coordinates are frequently omitted from final models despite their biological importance. To address these omissions, we have developed a new framework for De novo Ribonucleoprotein modeling in Real-space through Assembly of Fragments Together with Electron density in Rosetta (DRRAFTER). We show that DRRAFTER recovers near-native models for a diverse benchmark set of small RNA-protein complexes, as well as for large RNA-protein machines, including the spliceosome, mitochondrial ribosome, and CRISPR-Cas9-sgRNA complexes where the availability of both high and low resolution maps enable rigorous tests. Blind tests on yeast U1 snRNP and spliceosomal P complex maps demonstrate that the method can successfully build RNA coordinates in real-world modeling scenarios. Additionally, to aid in final model interpretation, we present a method for reliable *in situ* estimation of DRRAFTER model accuracy. Finally, we apply this method to recently determined maps of telomerase, the HIV-1 reverse transcriptase initiation complex, and the packaged MS2 genome, demonstrating that DRRAFTER can be used to accelerate accurate model building in challenging cases.

## Introduction

Recent advances in cryo-electron microscopy (cryoEM) have led to new structural insights into many biologically important ribonucleoprotein (RNP) assemblies, including the spliceosome, ribosome, telomerase, and CRISPR complexes [1-4]. For the increasing number of these maps with regions of high-resolution density (<4.0 Å), it is possible to manually trace atomic coordinates to obtain full-atom models [5]. However, most high-resolution maps still contain regions of lower resolution in which manual coordinate tracing is not feasible [6, 7]. For these regions as well as for the sizable number of maps determined at lower resolution, atomic coordinates are often obtained by fitting known structures of smaller subcomponents into the density [8]. This procedure presents a particular challenge for RNA-protein assemblies, as it is typically difficult to experimentally determine the coordinates of RNA subcomponents in isolation. For this reason, RNA coordinates are frequently omitted from models of RNP complexes [9-12], highlighting the critical need for computational methods that can accurately build RNA coordinates *de novo* into density maps of RNP assemblies.

The majority of existing computational methods focus on protein model building and refinement [13-16]. These methods, many of which are based on well-established structure prediction algorithms, are able to build proteins *de novo* into both high-and lower-resolution maps, but at best can handle the presence of predetermined RNA structures [17]. In principle, RNA structure prediction algorithms [18] could be similarly adapted for modeling RNA coordinates *de novo* into cryoEM maps of RNPs, but these methods have not yet been expanded to model RNA-protein complexes. Tools capable of modeling RNA into density maps are therefore limited to automated coordinate tracing within high-resolution maps [19] and refinement of reasonable initial structures. Developed primarily for high-resolution crystallographic density maps, refinement tools such as ERRASER, PHENIX, RCrane, and RNABC can be used to improve the quality of RNA structures [20-23]. Molecular dynamics flexible fitting refines reasonable starting structures, which are often previously determined structures of alternative conformational states, into density maps ranging from low-to high-resolution and has been successfully applied to large RNP assemblies such as the ribosome to generate accurate atomic models of different functional states [24]. However, there are currently no tools that are capable of building RNA structures *de novo* into low-resolution density maps.

Here, we have developed a computational framework for **D**e novo **R**NP modeling in **R**eal-space through **A**ssembly of **F**ragments **T**ogether with **E**lectron density in **R**osetta (DRRAFTER). We benchmarked DRRAFTER on pairs of high-(≤3.7 Å) and lower-resolution density maps for ten small RNA-protein complexes, the mitochondrial ribosome, spliceosomal tri-snRNP, and CRISPR Cas9-sgRNA complexes, and performed additional blind tests on maps of the yeast U1 snRNP and spliceosomal P-complex. These tests show that the accuracy of DRRAFTER models is comparable to that of models built by individually fitting subcomponent crystal structures, and importantly, that DRRAFTER model accuracy can be reliably estimated *in silico*. Additionally, application of our method to the recently determined 8.9 Å and 8.0 Å resolution telomerase and HIV-1 reverse transcriptase initiation complex (RTIC) maps recovered models that agree within error with previously published manually built models while requiring significantly reduced human effort, demonstrating that DRRAFTER can be used to accelerate and reduce bias in model building for lower resolution maps of RNPs. Finally, we used DRRAFTER to build a full-atom model of 1508 resolved nucleotides of the packaged MS2 genome, which until now had not been possible.

## Results

An overview of the DRRAFTER framework is shown in Figure 1. Briefly, known structures of protein components as well as RNA helices should first be individually fit into a density map (Figure 1A, B). This step is manual but rapid. For map subregions with missing RNA coordinates (Figure 1C), full-atom models based on a user-supplied RNA secondary structure are automatically constructed within Rosetta through fragment-based RNA folding and docking (Figure 1D). During this stage, models are scored initially with the Rosetta low-resolution RNA-protein potential and finally with a full-atom energy function. Both energy functions account for RNA-RNA and RNA-protein interactions and are also supplemented with a score term that monitors agreement with the density map. The best ten scoring models are then refined with the PHENIX-ERRASER pipeline to produce the final structures [20].

**Figure 1.**
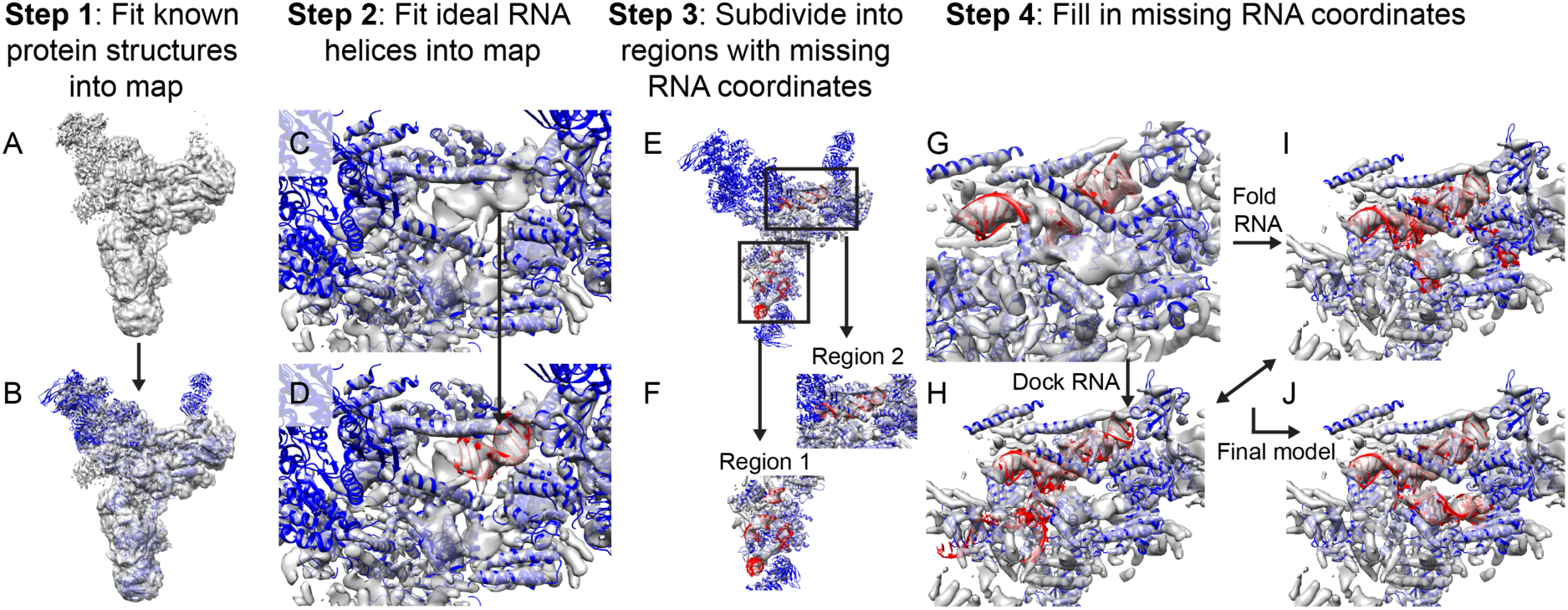
Overview of the DRRAFTER pipeline. (A) Starting from a cryoEM density map (here the 5.9 Å spliceosomal tri-snRNP map [9], gray), (B) individual protein structures (blue) are first fit into the density (here using Chimera). (C and D) Ideal RNA helices are then fit into the density map (red). (E) Subregions around the RNA helices where RNA coordinates are missing are visually identified, (F) and for each subregion, surrounding proteins and RNA helices are extracted from the larger model. (G) Each of these sub-structures is input into the DRRAFTER protocol in Rosetta, during which RNA coordinates are filled in through a Monte Carlo simulation involving (H) docking moves to optimize rigid body orientations within the density map and (I) RNA fragment insertions to fold the RNA (RNA coordinates colored red). Models are scored initially with a low-resolution RNA-protein energy function, which accounts for RNA-RNA and RNA-protein interactions, and finally by an all-atom potential, each supplemented with a score term that rewards agreement with the density map to produce (J) final models that fit into the density map.

To determine the accuracy of the method, we benchmarked DRRAFTER on RNA-protein systems with pairs of density maps at high-(≤3.7 Å) and lower-resolution (4.5-7 Å overall; 5.0-9.8 Å local resolution). Examples of the high-and lower-resolution density maps are shown in Figure S1. The benchmark set included ten small RNA-protein crystal structures for which we simulated density maps at both 5.0 and 7.0 Å resolution (Figure S2) [25-34], in addition to three large RNP machines with published experimental density maps containing regions where RNA coordinates had not previously been modeled: the spliceosomal tri-snRNP [9, 35], the CRISPR-Cas9-sgRNA complex [36, 37], and the mitoribosome [10, 38] (Figure 2). These systems represent a diverse range of RNA and RNA-protein structures including complex RNA junctions and interactions between proteins and both single-stranded and highly structured RNAs. To first establish the baseline target accuracy, we compared coordinates from the three lower-resolution experimental maps for the protein regions (for all three systems) and RNA regions (for the mitoribosome only) that were modeled into those maps to the later determined high-resolution coordinates. The root mean square deviations (RMSD) ranged from 1.3-9.1 Å (Figure 2A; see Methods). We then used DRRAFTER to build models of the ten small RNA-protein systems using the 5 and 7 Å simulated density maps, as well as six regions of the three large RNP machines using the lower-resolution experimental maps (local resolutions varied from 5.0-9.8 Å). Qualitatively, the DRRAFTER models closely recapitulate the overall folds of the high-resolution coordinates in all cases (Figure S2, Figure 2B-D). The RMSD accuracy of DRRAFTER models ranges from 0.7 Å to 6.2 Å (best of ten models, median of ten models was similar; Table 1), which is well within our targeted baseline accuracy range (Figure 2A). Additionally, the real-space correlation coefficients of the RNA models are comparable to the correlation of the high-resolution coordinates to the lower-resolution map (Table S1, Figure S3).

**Table 1.**
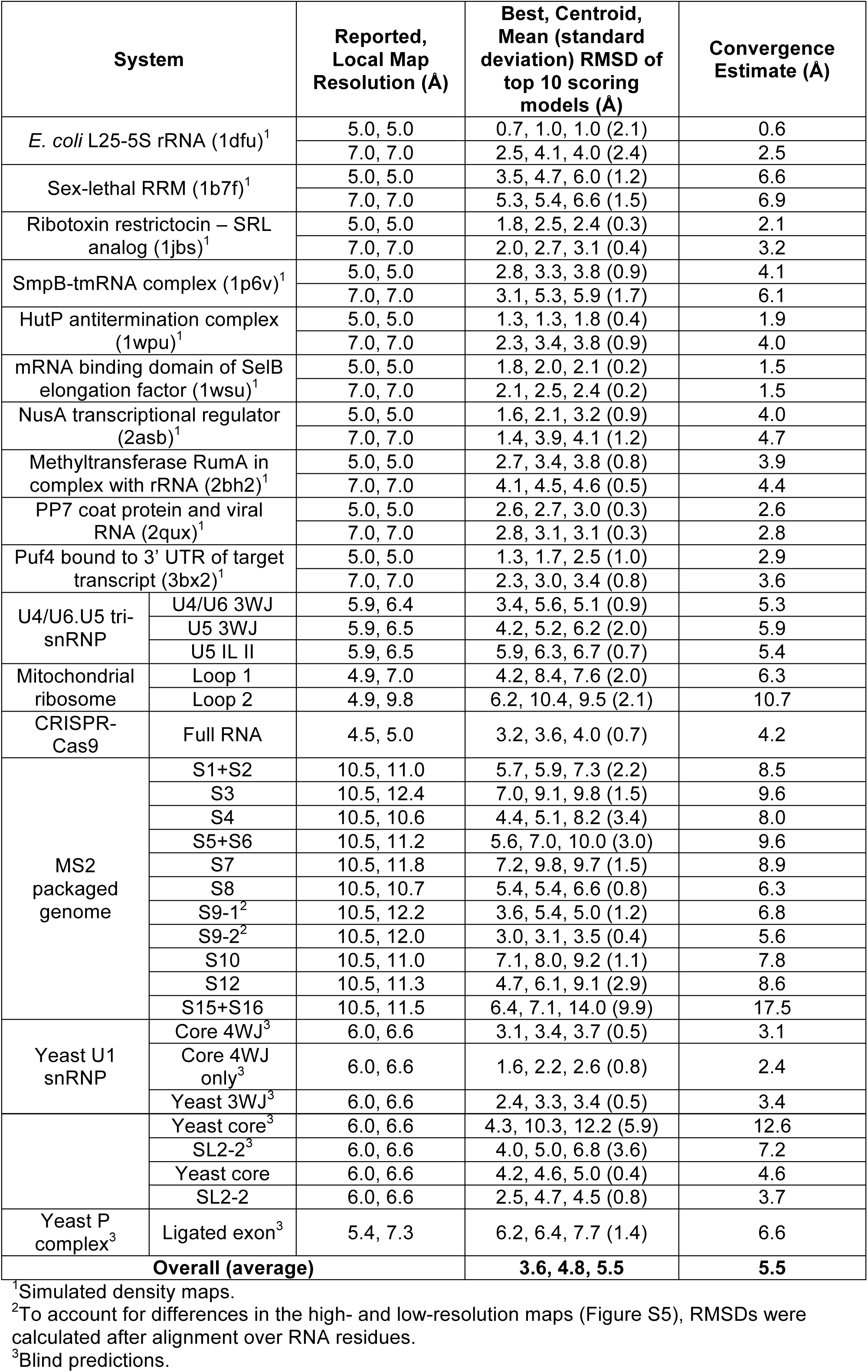
Local resolution and RMSD accuracy for all systems

| System |  | Reported,<br>Local Map<br>Resolution (Å) | Best, Centroid,<br>Mean (standard<br>deviation) RMSD of<br>top 10 scoring<br>models (Å) | Convergence<br>Estimate (Å) |
| --- | --- | --- | --- | --- |
| *E. coli* L25-5S rRNA (1dfu)<sup>1</sup> |  | 5.0, 5.0 | 0.7, 1.0, 1.0 (2.1) | 0.6 |
|  |  | 7.0, 7.0 | 2.5, 4.1, 4.0 (2.4) | 2.5 |
| Sex-lethal RRM (1b7f)<sup>1</sup> |  | 5.0, 5.0 | 3.5, 4.7, 6.0 (1.2) | 6.6 |
|  |  | 7.0, 7.0 | 5.3, 5.4, 6.6 (1.5) | 6.9 |
| Ribotoxin restrictocin – SRL<br>analog (1jbs)<sup>1</sup> |  | 5.0, 5.0 | 1.8, 2.5, 2.4 (0.3) | 2.1 |
|  |  | 7.0, 7.0 | 2.0, 2.7, 3.1 (0.4) | 3.2 |
| SmpB-tmRNA complex (1p6v)<sup>1</sup> |  | 5.0, 5.0 | 2.8, 3.3, 3.8 (0.9) | 4.1 |
|  |  | 7.0, 7.0 | 3.1, 5.3, 5.9 (1.7) | 6.1 |
| HutP antitermination complex<br>(1wpu)<sup>1</sup> |  | 5.0, 5.0 | 1.3, 1.3, 1.8 (0.4) | 1.9 |
|  |  | 7.0, 7.0 | 2.3, 3.4, 3.8 (0.9) | 4.0 |
| mRNA binding domain of SelB<br>elongation factor (1wsu)<sup>1</sup> |  | 5.0, 5.0 | 1.8, 2.0, 2.1 (0.2) | 1.5 |
|  |  | 7.0, 7.0 | 2.1, 2.5, 2.4 (0.2) | 1.5 |
| NusA transcriptional regulator<br>(2asb)<sup>1</sup> |  | 5.0, 5.0 | 1.6, 2.1, 3.2 (0.9) | 4.0 |
|  |  | 7.0, 7.0 | 1.4, 3.9, 4.1 (1.2) | 4.7 |
| Methyltransferase RumA in<br>complex with rRNA (2bh2)<sup>1</sup> |  | 5.0, 5.0 | 2.7, 3.4, 3.8 (0.8) | 3.9 |
|  |  | 7.0, 7.0 | 4.1, 4.5, 4.6 (0.5) | 4.4 |
| PP7 coat protein and viral<br>RNA (2qux)<sup>1</sup> |  | 5.0, 5.0 | 2.6, 2.7, 3.0 (0.3) | 2.6 |
|  |  | 7.0, 7.0 | 2.8, 3.1, 3.1 (0.3) | 2.8 |
| Puf4 bound to 3' UTR of target<br>transcript (3bx2)<sup>1</sup> |  | 5.0, 5.0 | 1.3, 1.7, 2.5 (1.0) | 2.9 |
|  |  | 7.0, 7.0 | 2.3, 3.0, 3.4 (0.8) | 3.6 |
| U4/U6.U5 tri-snRNP | U4/U6 3WJ | 5.9, 6.4 | 3.4, 5.6, 5.1 (0.9) | 5.3 |
|  | U5 3WJ | 5.9, 6.5 | 4.2, 5.2, 6.2 (2.0) | 5.9 |
|  | U5 IL II | 5.9, 6.5 | 5.9, 6.3, 6.7 (0.7) | 5.4 |
| Mitochondrial<br>ribosome | Loop 1 | 4.9, 7.0 | 4.2, 8.4, 7.6 (2.0) | 6.3 |
|  | Loop 2 | 4.9, 9.8 | 6.2, 10.4, 9.5 (2.1) | 10.7 |
| CRISPR-Cas9 | Full RNA | 4.5, 5.0 | 3.2, 3.6, 4.0 (0.7) | 4.2 |
| MS2<br>packaged<br>genome | S1+S2 | 10.5, 11.0 | 5.7, 5.9, 7.3 (2.2) | 8.5 |
|  | S3 | 10.5, 12.4 | 7.0, 9.1, 9.8 (1.5) | 9.6 |
|  | S4 | 10.5, 10.6 | 4.4, 5.1, 8.2 (3.4) | 8.0 |
|  | S5+S6 | 10.5, 11.2 | 5.6, 7.0, 10.0 (3.0) | 9.6 |
|  | S7 | 10.5, 11.8 | 7.2, 9.8, 9.7 (1.5) | 8.9 |
|  | S8 | 10.5, 10.7 | 5.4, 5.4, 6.6 (0.8) | 6.3 |
|  | S9-1<sup>2</sup> | 10.5, 12.2 | 3.6, 5.4, 5.0 (1.2) | 6.8 |
|  | S9-2<sup>2</sup> | 10.5, 12.0 | 3.0, 3.1, 3.5 (0.4) | 5.6 |
|  | S10 | 10.5, 11.0 | 7.1, 8.0, 9.2 (1.1) | 7.8 |
|  | S12 | 10.5, 11.3 | 4.7, 6.1, 9.1 (2.9) | 8.6 |
|  | S15+S16 | 10.5, 11.5 | 6.4, 7.1, 14.0 (9.9) | 17.5 |
| Yeast U1<br>snRNP | Core 4WJ<sup>3</sup> | 6.0, 6.6 | 3.1, 3.4, 3.7 (0.5) | 3.1 |
|  | Core 4WJ<br>only<sup>3</sup> | 6.0, 6.6 | 1.6, 2.2, 2.6 (0.8) | 2.4 |
|  | Yeast 3WJ<sup>3</sup> | 6.0, 6.6 | 2.4, 3.3, 3.4 (0.5) | 3.4 |
|  | Yeast core<sup>3</sup> | 6.0, 6.6 | 4.3, 10.3, 12.2 (5.9) | 12.6 |
|  | SL2-2<sup>3</sup> | 6.0, 6.6 | 4.0, 5.0, 6.8 (3.6) | 7.2 |
|  | Yeast core | 6.0, 6.6 | 4.2, 4.6, 5.0 (0.4) | 4.6 |
|  | SL2-2 | 6.0, 6.6 | 2.5, 4.7, 4.5 (0.8) | 3.7 |
| Yeast P complex<sup>3</sup> | Ligated exon<sup>3</sup> | 5.4, 7.3 | 6.2, 6.4, 7.7 (1.4) | 6.6 |
| **Overall (average)** |  |  | **3.6, 4.8, 5.5** | **5.5** |

<sup>1</sup>Simulated density maps.
<sup>2</sup>To account for differences in the high- and low-resolution maps (Figure S5), RMSDs were calculated after alignment over RNA residues.
<sup>3</sup>Blind predictions.

**Figure 2.**
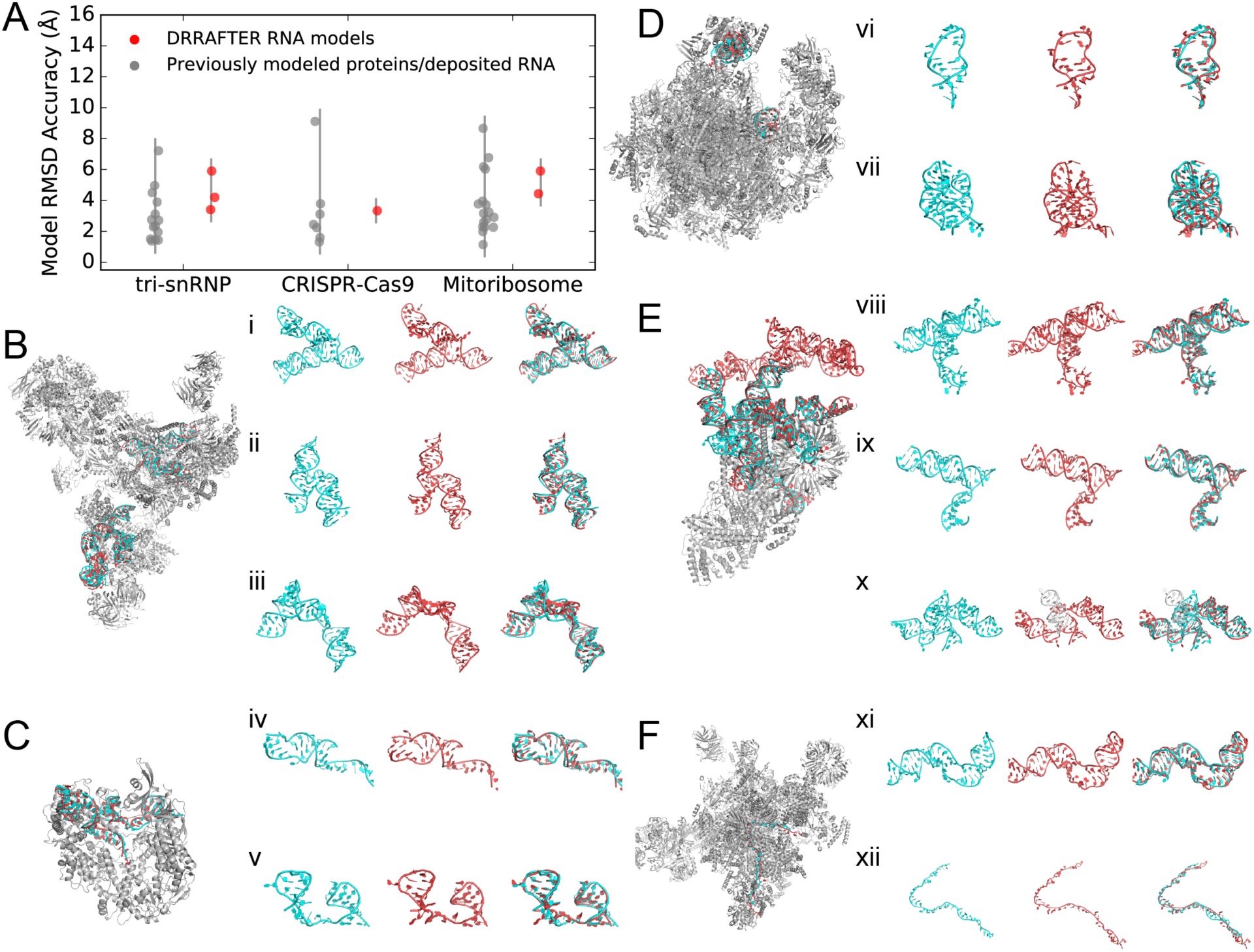
DRRAFTER recovers near-native models over a diverse benchmark set and two blind test cases. (A) RMSDs of DRRAFTER models (red; each region modeled is plotted as a separate point) and previously modeled low-resolution protein and RNA coordinates (gray; each protein or region of RNA is plotted as a separate point) compared with later determined high-resolution coordinates. (B-F) DRRAFTER models built into low-resolution maps (RNA colored red) overlaid with high-resolution coordinates (RNA colored cyan; protein colored silver) for (B) the spliceosomal tri-snRNP, (C) CRISPR-Cas9-sgRNA complex, (D) mitoribosome, (E) yeast U1 snRNP, and (F) yeast spliceosomal P complex. (i-xii) High-resolution RNA coordinates (left, cyan), RNA coordinates from DRRAFTER models built into low-resolution maps (middle, red), and high-resolution coordinates and DRRAFTER models overlaid (right) for the spliceosomal tri-snRNP (i) U4 / U6 three-way junction, (ii) U5 three-way junction, (iii) U5 internal loop II; CRISPR-Cas9-sgRNA complex (iv) sgRNA residues 11-30 and 57-68, (v) sgRNA residues 69-99; mitoribosome (vi) loop 1, (vii) loop 2; yeast U1 snRNP (blind) (viii) core four-way junction, (ix) yeast three-way junction, (x) yeast four-way junction (DRRAFTER model of SL3-2, SL3-3, and SL3-5 colored red; DRRAFTER model of SL3-4 colored white), (xi) SL2-2; yeast spliceosomal P complex (xii) ligated exon.

To test the applicability of DRRAFTER to higher resolution density maps, for each of the test cases in the benchmark set we also used DRRAFTER to build models into the high-resolution experimental density maps or, for the ten small crystal structures, simulated maps at 3 Å resolution (Figure S2). While the reported resolutions for the experimental maps were all better than 3.7 Å, the local resolution varied from 2.9 Å to 5.7 Å (Table S2). Compared to the published manually generated coordinates, the RMSDs of the DRRAFTER models ranged from 0.3 Å to 3.9 Å (Table S2), with the worst RMSD for the spliceosomal tri-snRNP U5 internal loop II (3.9 Å), which also had the lowest resolution density (5.7 Å). These results suggest that while the DRRAFTER framework is primarily intended for cases where manual coordinate tracing is not feasible, it can be used to automatically build coordinates into high-resolution maps, though in some cases final manual adjustments may be necessary and careful visual inspection is always recommended.

As a rigorous challenge, K.K. and R.D. performed blind tests of the DRRAFTER pipeline on early stage 6.0 Å and 5.4 Å resolution maps of the yeast U1 snRNP and spliceosomal P complex, respectively, prior to the publication of higher-resolution maps with resolutions of 3.6 Å and 3.3 Å, respectively (kept hidden by S.L., H.Z., R.Z.) [39, 40]. The yeast U1 snRNP modeling was carried out over a period of three days, during which we built DRRAFTER models of five subregions covering the majority of the 568-nucleotide U1 snRNA. A previously published structure of the core human U1 snRNP helped identify the location of the core four-way junction in the map, but because the human structure did not fit well in the density map and the yeast snRNA is significantly larger than the human U1 snRNA (568 vs. 164 nucleotides), nearly the entire RNA was modeled *de novo* (Figure 2E). Blind DRRAFTER models of the core four-way junction (Figure 2E (viii)) and yeast core three-way junction regions (Figure 2E (ix)) achieved RMSDs of 3.1 Å and 2.4 Å, respectively, with residues within the four-way junction reaching 1.6 Å RMSD accuracy. The best model of SL2-2 achieved RMSD accuracy of 4.0 Å (Figure 2E (xi)), although we noted that models of this region suffered from a lack of compute time (∽450 models generated vs. target of 3000 models). When later revisited with additional computational expenditure (∽3000 models generated), the RMSD dropped to 2.5 Å. The best model of the yeast core four-way junction over SL3-2, SL3-3, and SL3-5 achieved RMSD accuracy of 4.3 Å (Figure 2E (x)). SL3-4 was excluded from the final RMSD calculation because we were unable to build a model that fit into the density, as determined by visual inspection. After unblinding the high-resolution coordinates, we learned that the proposed secondary structure for this region, which was enforced during the DRRAFTER modeling, was incorrect. When this region was subsequently revisited with the corrected secondary structure, we were able to build models with SL3-4 in the density, and the RMSD accuracy over the entire yeast core four-way junction improved slightly to 4.2 Å. Finally, we could not assess the accuracy of models that we built for the peripheral SL3-7 domain because coordinates were not built into the final map, which only showed diffuse density for that region. We provide a complete all-atom model for the yeast U1 snRNP, including these peripheral regions, in Supporting Information (Supplemental File 1).

When modeling the yeast spliceosomal P complex, we discovered that the majority of the density could be modeled well by the previously published structure of the C* complex, the state immediately prior to P complex formation in the catalytic cycle of the spliceosome [41, 42]. We therefore focused our attention on the structure of the ligated exon, which is not yet present in the C* complex. This long single-stranded RNA region proved challenging to model as indicated by two measures. First, the density in this region was at 7.3 Å resolution, considerably poorer than the overall 5.4 Å resolution of the map. Second, our final pool of DRRAFTER models exhibited substantial structural heterogeneity. Indeed, while our models cluster around the high-resolution coordinates, the RMSD accuracy of our best model was 6.2 Å, poorer than for the majority of the test cases in our benchmark set (Figure 2F).

Inspired by the challenge of these blind tests, we sought to develop a method to estimate the accuracy of DRRAFTER models *in silico*. This would allow model quality to be quantitatively determined in realistic modeling scenarios. We identified two metrics that are predictive of final model accuracy. First, the local resolution places approximate bounds on the final modeling accuracy (Figure 3A). The correlation is significant (two-tailed p = 4×10^−8^ for Pearson’s correlation coefficient, N = 61, data listed in Table 1), but weak (R^2^ = 0.21) suggesting that there are additional factors that determine model accuracy. Second, we assessed the convergence of DRRAFTER models by calculating the average pairwise RMSD over the best ten scoring models (Table 1, Table S2, Figure S4). This convergence estimate is correlated with the accuracy of the best of the top ten models (Figure 3B, Table 1, R^2^ = 0.67, two-tailed p = 6×10^−16^, N = 61; excluding models with convergence > 12 Å: R^2^ = 0.78, two-tailed p = 3×10^−20^, N = 59), the centroid of the top ten models (Figure 3C, Table 1, R^2^ = 0.72, two-tailed p = 4×10^−18^, N = 61; excluding models with convergence > 12 Å: R^2^ = 0.82, two-tailed p = 2×10^−22^, N = 59), and the mean accuracy of the top ten models (Figure 3D, Table 1, R^2^ = 0.93, two-tailed p = 4×10^−36^, N = 61; excluding models with convergence > 12 Å: R^2^ = 0.92, two-tailed p = 1×10^−33^, N = 59). With these results we expect that prior to modeling, the local map resolution can be used to place bounds on the expected modeling accuracy, and after modeling is completed, the convergence of the DRRAFTER models can be used to reliably estimate modeling accuracy.

**Figure 3.**
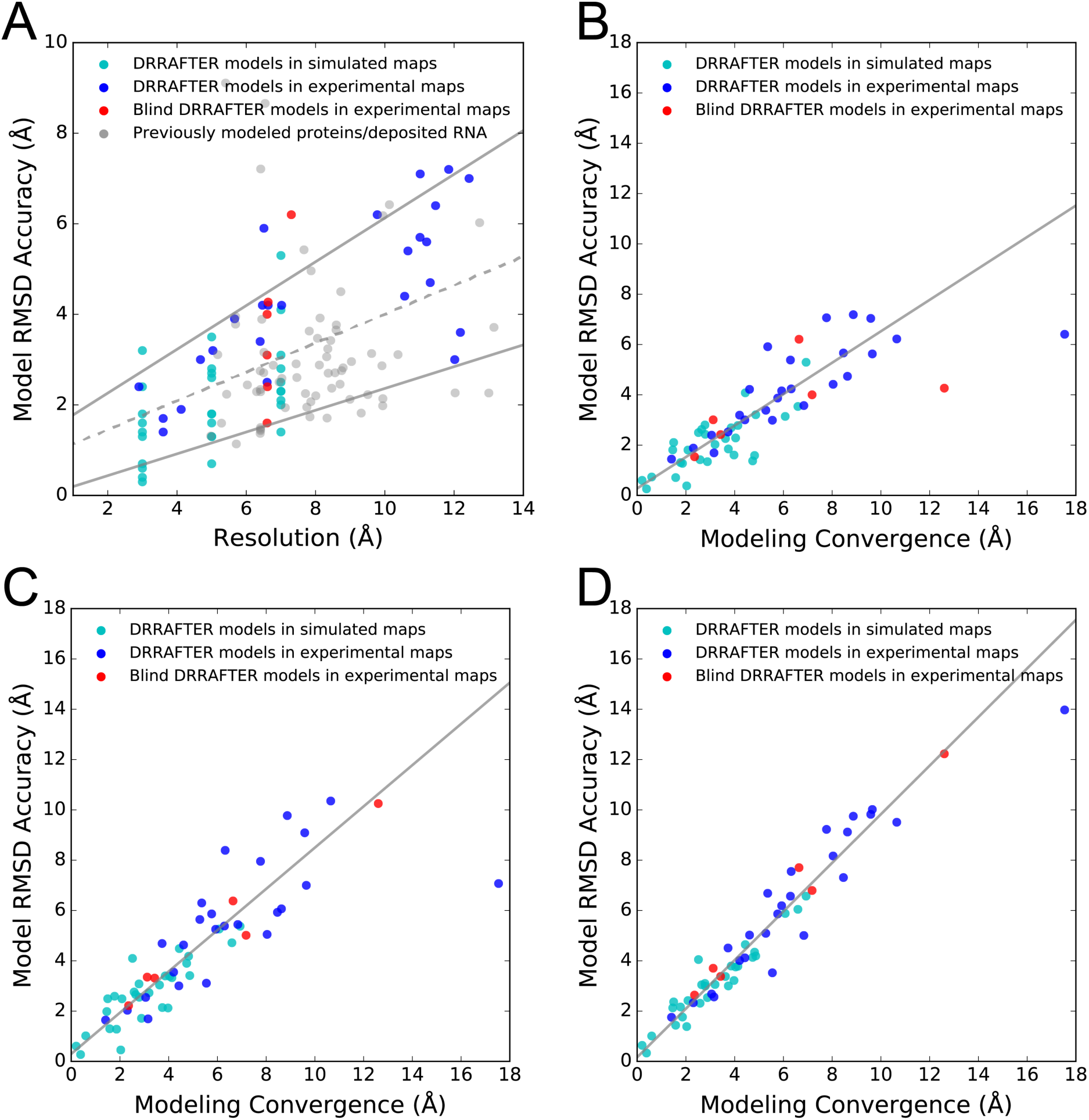
Estimating DRRAFTER model accuracy. (A) RMSD accuracy versus local map resolution (Table 1 and Table S2) for DRRAFTER models built into high-and low-resolution simulated (cyan) and experimental maps (blue), blind DRRAFTER models built into low-resolution experimental maps (red), and previously modeled low-resolution protein and RNA coordinates (gray). The best-fit line (dashed gray) is given by y = 0.32x + 0.81. The best-fit upper and lower bound lines (solid gray) are given by y = 0.48x + 1.29 and y = 0.24x-0.04, respectively (see Methods). RMSD accuracy versus DRRAFTER modeling convergence for (B) the most accurate of the top ten scoring DRRAFTER models, (C) the centroid of the top ten scoring DRRAFTER models, and (D) the mean RMSD to native across the top ten scoring DRRAFTER models. The best-fit lines (gray; excluding the two points with convergence > 12 Å: MS2 S15 + S16 and blind yeast U1 snRNP yeast core) are given by (B) y = 0.62x + 0.28, (C) y = 0.82x + 0.30, (D) y = 0.97x + 0.17.

For RNP targets of exceptional biological value, researchers have committed extraordinary efforts to manually piece together RNA models within low-resolution maps of RNPs. In the few cases where this manual model building is actually feasible, it is extremely time-consuming and subject to considerable bias. We therefore wanted to test whether DRRAFTER could be used to accelerate model building and reduce human bias in these cases. We applied DRRAFTER to the recently determined 8.9 Å map of *Tetrahymena* telomerase and the 8.0 Å map of the HIV-1 reverse transcriptase initiation complex (RTIC), where models of the RNA had previously been built manually [43, 44]. The DRRAFTER models agree well with the published models with mean RMSDs over the top ten models of 5.7 Å for HIV-1 RTIC and 7.6 Å for telomerase (6.6 Å excluding the poorly converged single stranded RNA residues 52-68) (Figure 4A, B, C, D). Building these models with DRRAFTER required only a few hours of human effort, versus the days to weeks that are usually required for manual model building. Additionally, by using DRRAFTER to build these models we were able to calculate their expected accuracy. Using the convergence of the DRRAFTER models, we estimate that the best of the ten DRRAFTER models have RMSD accuracies to the “true” coordinates of 3.5 Å for telomerase (convergence = 5.2 Å), and 4.2 Å (convergence = 6.3 Å) for the HIV-1 RTIC RNA.

**Figure 4.**
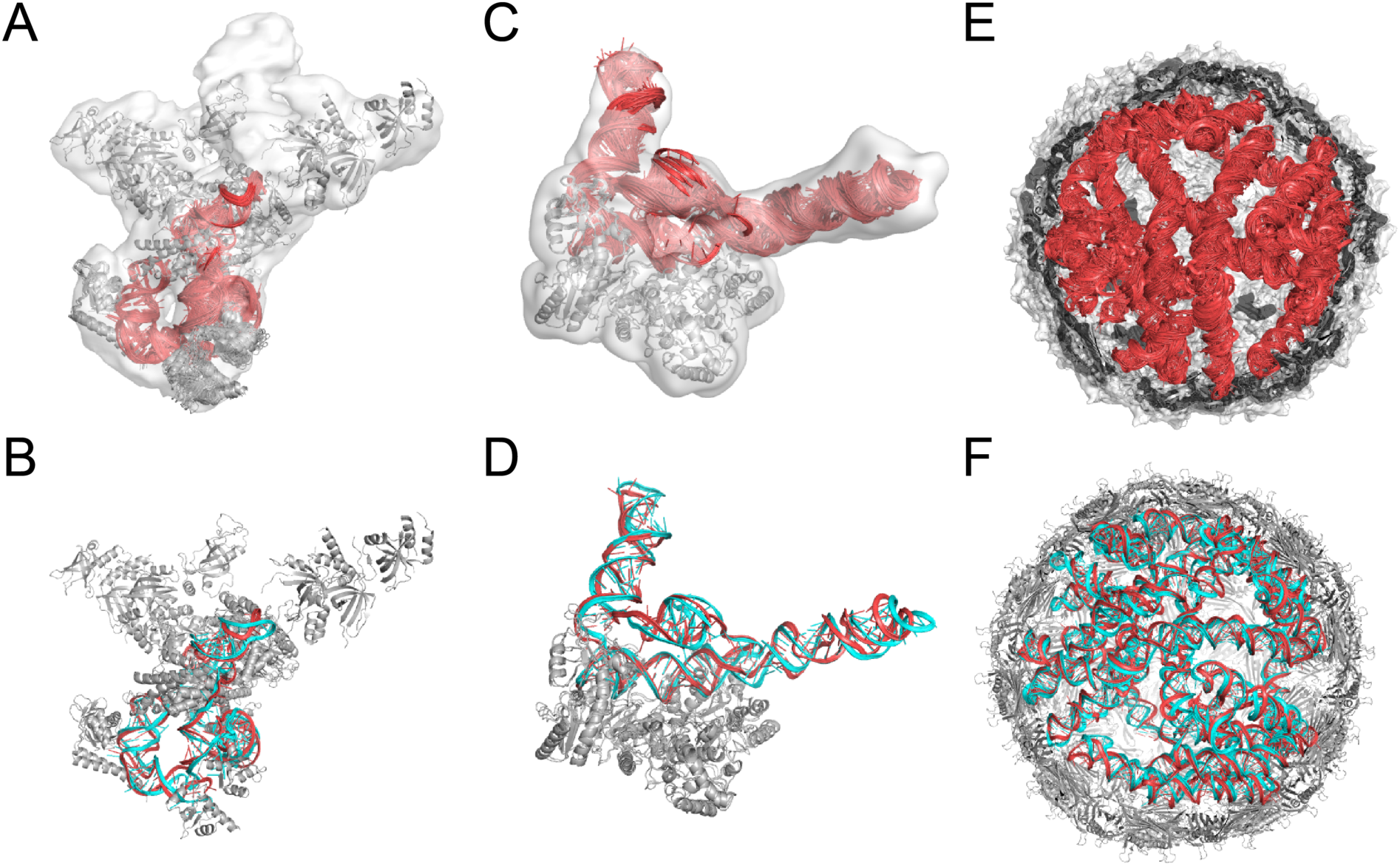
DRRAFTER can accelerate manual model building into low-resolution density maps. Overlay of ten best scoring DRRAFTER models (RNA colored red, protein colored gray) for (A) telomerase, (C) HIV-1 RTIC, and (E) the packaged MS2 genome (built into the 10.5 Å resolution map). Regions with more variability between models are estimated to be less accurate. Overlay of DRRAFTER models (RNA colored red, protein colored gray) with previously built manual models (RNA colored cyan, protein colored gray) for (B) telomerase and (D) HIV-1 RTIC. A single DRRAFTER model (centroid) of the ten best scoring is shown for clarity. (F) Overlay of DRRAFTER models built into independently determined 3.6 Å (cyan) and 10.5 Å (red) resolution maps of the packaged MS2 genome.

Finally, we applied DRRAFTER to the recently determined 3.6 Å map of the packaged MS2 genome [45]. Despite the high-resolution overall, the local resolution in the region of the packaged RNA was not high enough for a full-atom model to be built, with the exception of several protein-bound RNA hairpins. With DRRAFTER, we were able to build a model of 1508 nucleotides (Figure 4; Supplemental File 2) with estimated accuracies of 2.4-6.0 Å (convergence = 3.8-9.7 Å). As a final test of DRRAFTER accuracy, we additionally applied the framework to the previously published 10.5 Å map of the packaged MS2 genome and compared the resulting models to those based on the 3.6 Å map [46]. The RMSDs are between 3.0 Å and 7.2 Å; qualitatively, the models agree very well, and many of the differences in the models reflect underlying differences in the 3.6 Å and 10.5 Å maps (Figure S5).

## Discussion

For systems representing all major classes of RNPs with maps of a wide range of resolutions, DRRAFTER was able to successfully build near-native coordinates in regions where manual coordinate tracing was difficult or intractable. Over a benchmark set of both simulated and experimental maps, DRRAFTER models consistently recovered native RNA folds. Separate blind tests of the method demonstrate that the DRRAFTER framework can be successfully applied in realistic modeling settings. Additionally, even in cases where manual modeling into low-resolution maps may be feasible, it is slow, painstaking, and can suffer from errors; DRRAFTER can be used to accelerate and reduce bias from the process. DRRAFTER has the added advantage over manual modeling of providing a way to estimate model accuracy, which should aid in interpretation of final models. Overall, we expect that DRRAFTER will be widely useful for building RNA coordinates into cryoEM maps.

The tests presented here suggest three main areas for future improvement of the DRRAFTER pipeline. First, DRRAFTER relies on having accurate RNA secondary structure information. In some cases, the current DRRAFTER pipeline may be able to distinguish between different secondary structure possibilities; for the U1 snRNP yeast core four-way junction test case, models with the incorrect secondary structure were unable to fit into the density, while later models with the corrected secondary structure fit well. However, this strategy is unlikely to be feasible in cases where large sections of an RNA secondary structure are unknown and/or the number of possible secondary structures is large. We expect that combining cryoEM data and the DRRAFTER pipeline with NMR or biochemical techniques that probe RNA secondary structure will be critical to solving accurate structures for many RNPs [47].

Second, improvement to the final accuracy of DRRAFTER models will require advances in structure refinement tools. Existing refinement methods such as the PHENIX-ERRASER pipeline used here work best with high-resolution density maps and near atomic accuracy starting models. DRRAFTER model refinement will benefit from new tools that can handle more substantial structural changes and focus on refinement into lower-resolution maps.

Third, DRRAFTER does not remodel protein backbones or build missing protein coordinates. DRRAFTER may therefore build RNA coordinates into nearby unfilled protein density. This challenge can often be overcome by segmenting out density that is visually recognizable as belonging to a protein prior to DRRAFTER modeling. However, in some cases it is difficult to distinguish between density belonging to proteins and RNA. It may also be more challenging to sample the correct protein-bound RNA conformation when the protein partner is not present. Ultimately, integrating DRRAFTER with existing protein structure modeling tools will be necessary to complete the pipeline for RNP model building.

Lastly, DRRAFTER automates RNA model building and error estimation, but final visual inspection should still play an important role in the modeling process. We recommend visually inspecting at least the top ten DRRAFTER models; a similar process has been powerful for our ERRASER tool [48, 49]. Particularly when the modeling error is predicted to be high, visual examination can identify regions for which modeling assumptions, such as the secondary structure or initial placements of proteins and RNA helices, may be incorrect.

## Acknowledgements

We thank members of the Das lab for useful discussions and members of the Rosetta community for discussions and code sharing. Calculations were performed on the Stanford BioX^3^ cluster, supported by NIH Shared Instrumentation Grant 1S10RR02664701. This work was supported by a Gabilan Stanford Graduate Fellowship (K.K.), a NSF GRFP (K.K.), T32-GM008294 (Molecular Biophysics Training Program; K.P.L. and K.K), NIGMS MIRA R35 GM122579 (R.D.), R01 GM121487 (R.D.), and NIH grant GM114178 (R.Z.).

## Author Contributions

K.K. and R.D. designed the computational approach. K.K. implemented the method and performed the tests and analysis. S.L., Z.H.Z., and R.Z. provided the U1 snRNP and P complex blind test cases. K.P.L., G.S., E.V.P, and J.D.P. provided the HIV-1 RTIC test case and provided initial feedback on the method. K.K. and R.D. wrote the manuscript with input from S.L., K.P.L., G.S., E.V.P., J.D.P., Z.H.Z., and R.Z.

## Methods

### The DRRAFTER pipeline

For each system, all available structures of individual proteins were collected from the PDB and then fit into the cryoEM density map in Chimera using the “Fit in Map” function [50]. Ideal A-form RNA helices were built with the Rosetta tool, rna_helix.py, and then fit into the maps in Chimera [50]. Following conventional protocols [9-12], these steps were performed manually, but completed rapidly (minutes per structure). Regions with missing RNA coordinates were identified and subdivided by visual inspection. The surrounding RNA helices and proteins were extracted from the overall model of the RNP and used as the input to the Rosetta DRRAFTER run.

The Rosetta stage consists of a modified version of the FARFAR method, run through the Rosetta rna_denovo application [51, 52]. The method was updated so that both proteins and density maps can be included. There are two stages to this protocol. First, a low resolution Monte Carlo stage, which includes standard RNA fragment insertion moves to fold the RNA, now allows docking moves that optimize the placement of RNA helices and proteins. Docking moves for RNA helices include rotations and translations about the helical axis, in addition to the standard random rigid body perturbations. During this stage, the proteins are treated as rigid bodies. Each conformation is scored with the low-resolution RNA-protein potential in Rosetta [53], augmented by the “elec_dens_fast” score term, which scores the agreement between the map and model [54].

After the low-resolution stage, the structure goes through full-atom refinement. First, the structure is subjected to energy minimization in which the RNA as well as the protein sidechains within a 20.0 Å distance of any RNA atom are allowed to move. Then, the structure is further refined through single residue fragment insertions, sidechain packing, and small rigid body perturbations. The structure is then subjected to a second round of energy minimization. Scoring during these phases is performed with the full-atom Rosetta energy function, which includes terms that describe hydrogen bonding, electrostatics, torsional energy, van der Waals interactions and solvation, and is also supplemented with the density score term elec_dens_fast [54, 55]. This score function is available within Rosetta as “rna_hires_with_protein.wts”. The top ten models are output from the run, with the centroid model highlighted, to be visually inspected and to allow final manual selection. The DRRAFTER code will be freely available to academic users as part of the Rosetta software package in releases after March 14, 2018 (www.rosettacommons.org) and is automatically compiled along with ERRASER, which is already in routine use for RNA and RNP cryoEM.

An example Rosetta command line is as follows:

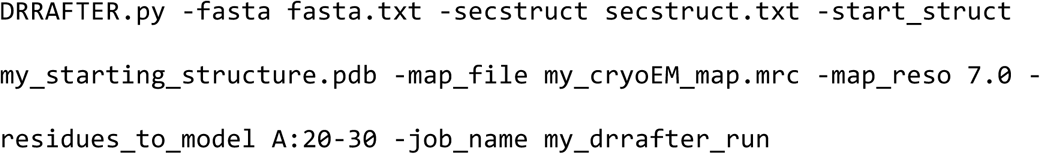

where fasta.txt is a FASTA file listing the full sequence of the complex, secstruct.txt is a file containing the secondary structure in dot bracket notation (with dots for protein residues),-residues_to_model (here given a value of A:20-30) specifies the residues that should be built in the DRRAFTER run, my_starting_structure.pdb is the PDB file containing all fit protein structures and RNA helices,-map_file specifies the density map,-map_reso specifies the resolution of the map, and-job_name specifies a name for the run (which controls the names of the output files). Documentation and a demo are available at www.rosettacommons.org.

Modeling convergence was calculated by taking the average of the pairwise RMSDs over the RNA region being modeled for the best ten scoring DRRAFTER models. An example command line to calculate convergence and corresponding error estimates is as follows:

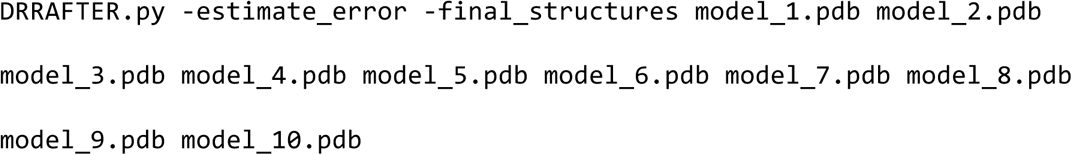

Approximately 3000 DRRAFTER models were generated in all cases, and the top ten scoring were then subjected to the PHENIX-ERRASER pipeline [20]. For the PHENIX runs, secondary structure restraints were automatically generated using phenix.secondary_structure_restraints and applied during refinement with phenix.real_space_refine. Additionally, coordinate restraints were applied for all residues in RNA helices. During the ERRASER runs, the first base pair of each RNA helix was kept fixed, as well as any residues contacting a protein surface, or near enough that ERRASER introduced protein-RNA clashes if the residue was not kept fixed.

### Model analysis

RMSDs (reported in Table 1) were calculated over RNA heavy atoms after initial alignment over protein heavy atoms. These calculations were carried out in Rosetta and Pymol. RMSDs for previously modeled coordinates in the spliceosomal tri-snRNP were calculated for protein structures that had been fit into the lower-resolution (5.9 Å) density map in Chimera following the description in the methods section of the original paper [9] versus the high-resolution coordinates of the corresponding proteins in PDB ID 5GAN [35]. Homologous protein structures that were docked into the lower-resolution map were omitted from this calculation. For the mitoribosome, RMSDs were calculated between the coordinates deposited with the lower-resolution (4.9 Å) map (PDB ID: 4CE4) and the high-resolution (3.4 Å) map (PDB ID: 4V1A and 4V19) for proteins present in both as well as for RNA regions that could not have been modeled by simple threading of the *E. coli* ribosome structure. For the Cas9-sgRNA complex, the protein coordinates were taken from the crystal structure of CRISPR-Cas9 in complex with sgRNA and double stranded DNA (PDB ID 5F9R) and broken up into domains, and each of these was individually fit into the cryoEM density map [36]. RMSDs between these regions and the high-resolution crystal structure (PDB ID 4ZT0) were calculated.

Local map resolution was calculated with Resmap [7], then loaded into Chimera version 1.11.2 along with the corresponding high-resolution coordinates. The “Values at Atom Positions” tool in Chimera was used to find the local resolution at the positions of each of the atoms in the high-resolution structure. The values at the positions of all of the RNA atoms for the region being modeled were averaged (with a python script) to give the local resolution for that region.

Best-fit lines describing the upper and lower bounds of DRRAFTER model accuracy versus local resolution (Figure 3A) were calculated using the minimum RMSD values (lower bound) or 90^th^ percentile RMSD values (upper bound) in each 1 Å bin ranging from 2.5 to 12.5 Å local resolution.

Real-space correlation coefficients were calculated for RNA coordinates being modeled only (surrounding proteins were not included to facilitate comparison between high-and low-resolution coordinates) using the PHENIX tool phenix.get_cc_mtz_pdb with fix_xyz = True and scale = True. The “Map correlation in region of model” was reported.

Figures were generated with Pymol and UCSF Chimera.

### Simulated benchmark

Ten systems were chosen from the nonredundant set of RNA-protein complexes with corresponding unbound protein structures available, described in [56]. The specific systems were selected manually to represent a diversity of types of RNA-protein interactions (unbound protein structures listed in parentheses): 1DFU (1B75) [25, 57], 1B7F (3SXL) [26, 58], 1JBS (1AQZ) [27, 59], 1P6V (1K8H) [28, 60], 1WPU (1WPV) [29], 1WSU (1LVA) [30, 61], 2ASB (1K0R) [31, 62], 2BH2 (1UWV) [32, 63], 2QUX (2QUD) [33], and 3BX2 (3BWT) [34]. For each of these systems, density maps were simulated at 3.0 Å, 5.0 Å, and 7.0 Å resolution with the pdb2vol tool in the Situs package [64]. Unbound protein structures (listed above) were fit into the simulated density maps using Chimera’s Fit in Map tool. Ideal RNA helices for helical segments of RNA were generated with rna_helix.py in Rosetta and then fit into the maps using Chimera’s Fit in Map tool. For systems that contained only single-stranded RNA, an ideal A-form nucleotide was fit approximately into the map – throughout the later DRRAFTER simulation, it was allowed to change its conformation and orientation within the map. The remaining RNA residues were also built with the DRRAFTER protocol in Rosetta. The full protein structures were included in the simulations, and were allowed to dock as rigid bodies within the density map. The ideal RNA helices were also subjected to docking within the map to optimize their final placement.

### Spliceosomal tri-snRNP modeling

All proteins listed in Extended Data Table 1 of the original paper [9] were fit into the full tri-snRNP density map (EMD 2966), as well as the structure of the C-terminal fragment of PRP3, which had since been solved (PDB ID: 4YHU) [65]. Ideal RNA helices were fit into the map for all helical parts of the three regions modeled: the U5 snRNA three-way junction (residues 35-53, 62-91, and 103-119), the U5 snRNA internal loop II (residues 4-40, 114-144), and the U4 / U6 snRNA three-way junction consisting of U4 snRNA residues 1-64 and U6 snRNA residues 55-80. All RNA helices were allowed to move as rigid bodies throughout the DRRAFTER runs. Proteins were kept fixed. In each case, the density map was approximately segmented around the region of interest with the Segment Map tool in Chimera (Segger v1.9.4). RMSDs were calculated relative to the coordinates from the 3.7 Å map, PDB ID 5GAN [35].

For DRRAFTER models built into the 3.7 Å map (EMD 8012), the protein structures were taken from the corresponding PDB entry, 5GAN. Ideal RNA helices were fit into the map and DRRAFTER runs were performed as described above.

### Mitoribosome modeling

DRRAFTER models were built extending from the coordinates deposited with the 4.9 Å map (EMD 2490), PDB ID 4CE4 for two regions for which RNA coordinates were missing [10]. “Loop 1” consisted of RNA residues 401-407, and “Loop 2” consisted of RNA residues 495-547. Connected RNA residues were included in the simulations. For Loop 2, an initial model of residues 502-522 and 529-544 was built by taking H43 and H44 from the *E. coli* ribosome structure (PDB ID 4YBB) and threading in the mitoribosome sequence [66]. This model was fit approximately into the density map with the Fit in Map function in Chimera and then included as a rigid body, allowed to rotate and translate, in the DRRAFTER run. Models were similarly built into the 3.4 Å map (EMD 2787) [38], but surrounding protein and RNA coordinates were taken from PDB structures 4V19 and 4V1A (deposited with the 3.4 Å map).

### CRISPR-Cas9-sgRNA modeling

Protein coordinates were taken from the crystal structure of CRISPR-Cas9 in complex with sgRNA and double stranded DNA (PDB ID 5F9R) [36]. The protein was split up into seven domains (Arg, CTD, HNH, Helical-I, Helical-II, Helical-III, and RuvC) and each was fit individually into the 4.5 Å cryoEM map (EMD 3276) [36]. The protein domains were kept fixed throughout the DRRAFTER run. Ideal A-form RNA helices were fit into the map for all helical sections of the sgRNA. Models were built for sgRNA residues 11-99, but RMSDs were only computed over residues with coordinates in the 2.9 Å crystal structure, PDB 4ZT0 (residues 11-30 and 57-99) [37]. Models were similarly built into the 2.9 Å crystallographic density map (4ZT0), but with the protein coordinates taken from 4ZT0.

### Blind yeast U1 snRNP modeling

Modeling was performed with a 6.0 Å resolution map of the yeast U1 snRNP from an earlier stage of processing than the later published 3.6 Å map [39]. The core four-way junction region of the map was identified by fitting the structures of the human U1 snRNP (3CW1 and 3PGW) into the map [67, 68]. Structures of the seven yeast Sm proteins (B, D1, D2, D3, E, F, and G) were taken from PDB ID 5GMK and fit into the map with the Fit in Map tool in Chimera [69]. Homology models of PRP39 and PRP42 were generated with Modeller and fit into the map [70]. A homology model of the U1-70K RRM was fit into the map and later allowed to move as a rigid body. We assumed that the U1 snRNA would adopt the secondary structure proposed in the literature [71]. Ideal RNA helices were fit into the map for all helical regions of the RNA. DRRAFTER models were built for five regions of the RNA: the core four-way junction (residues 11-60, 154-178, and 534-559), the SL3-1/SL3-2/SL3-6 yeast three-way junction (residues 172-185, 304-325, and 526-539), the yeast core four-way junction (residues 181-202 and 236-308), and SL3-7 (residues 310-531).

### Blind yeast spliceosomal P complex modeling

Models of the P complex ligated exon were built into a 5.4 Å resolution map from an earlier stage of processing than the later published 3.3 Å map [40]. Previously determined structures of the yeast spliceosomal C* complex were fit into the map (PDB 5MQ0 and 5WSG), which allowed identification of the density for the ligated exon [41, 42]. Coordinates for PRP22 were taken from the C* complex (5MQ0) and fit into the density map individually. The coordinates of the RNA bound to PRP22 were modeled by taking the structure of PRP43 in complex with RNA (PDB ID 5I8Q) and aligning it to PRP22, then taking the resulting RNA coordinates from the complex [72]. These RNA coordinates were kept fixed relative to PRP22 in all DRRAFTER runs. DRRAFTER runs were set up with varying numbers of nucleotides spanning the exon-exon junction and the active site in PRP22, ranging from ten to twenty nucleotides. Models were selected from the runs with the fewest number of nucleotides spanning the exon-exon junction and the PRP22 active site in which there were no breaks in the RNA chain (thirteen and fourteen nucleotides).

### HIV-1 RTIC modeling

Approximate initial locations for all helical segments of the HIV-1 RNA and bound tRNA were determined by fitting ideal A-form helices into an 8.0 Å map of the HIV-1 RTIC [44]. The alternative tRNA secondary structure was assumed as in the previously published manual modeling. Protein coordinates were taken from the previously published model. Final refinement was carried out only with PHENIX, as was carried out for the previously published model. The fifteen best scoring models were visually inspected and the top ten without large distortions in the PBS helix were selected as the final set of ten best scoring models.

### Tetrahymena telomerase modeling

All proteins described in the original paper [43] were fit into the 8.9 Å map of *Tetrahymena* telomerase (EMD 6443). Additionally, the RNA pseudoknot (5KMZ) [43], RNA residues 155-159 bound to the N-terminal domain of the human La protein (2VOP) [73], the structure of the RNA TBE bound to the TRBD (5C9H) [74], and the RNA stem IV loop (2M21) [75], RNA stem IV (4ERD) [76], “half” an ideal A-form helix for the template RNA, and ideal A-form helices for the remaining helical regions of the RNA were fit into the map with Fit in Map in Chimera and then each allowed to move individually in the subsequent DRRAFTER runs. The full RNA was modeled as a single region.

### MS2 packaged genome modeling

The packaged MS2 genome was modeled based on the 3.6 Å map (EMD 8397) using the published proposed secondary structure [45]. Because the RNA density in this map is noisy, a 1.5 Å Gaussian filter was applied to the map in Chimera prior to RNA modeling (similarly, RNA density in the original paper [45] was examined after low-pass filtering to 6 Å resolution). Models were built for 10 regions: S1 + S2 (residues 29-227, 341-369); S3 (residues 372-583); S4 (residues 888-943); S5 + S6 (residues 963-1119); S7 (residues 1132-1283); S8 (residues 1714-1806); S9-1 (residues 1837-1896); S9-2 (residues 1900-1940); S10 (residues 1960-2122); S12 (residues 1810-1826, 2202-2340); S15 + S16 (residues 2346-2353, 2757-2661, 3088-3111, 3249-3382). The published coordinates for the protein capsid and bound RNA hairpins were kept fixed (5TC1) [45]. One ideal RNA helix for each region was fit into the map; the initial coordinates of the remaining helices were not provided for the DRRAFTER run (and were therefore determined by the initial random perturbations to the RNA structure). For comparison, models were similarly built into the 10.5 Å map (EMD 3403) [46], without the high-resolution coordinates of the RNA hairpins. Because the 3.6 Å and 10.5 Å maps differed significantly in regions S9-1 and S9-2 (Figure S5), RMSDs for these regions were calculated after alignment over all RNA heavy atoms. For all other regions, RMSDs were calculated over RNA heavy atoms after alignment over all protein residues.

## Data availability statement

The accession codes used in this study are as follows: *E. coli* L25-5S rRNA (PDB 1DFU and 1B75), sex-lethal RRM (PDB 1B7F and 3SXL), ribotoxin restrictocin sarcin-ricin loop analog (PDB 1JBS and 1AQZ), SmpB-tmRNA complex (PDB 1P6V and 1K8H), HutP antitermination complex (PDB 1WPU and 1WPV), mRNA binding domain of SelB elongation factor (PDB 1WSU and 1LVA), NusA transcriptional regulator (PDB 2ASB and 1K0R), methyltransferase RumA in complex with rRNA (PDB 2BH2 and 1UWV), PP7 coat protein and viral RNA (PDB 2QUX and 2QUD), Puf4 bound to 3’ UTR of target transcript (PDB 3BX2 and 3BWT), tri-snRNP (EMD 2966 and 8012; PDB 4YHU and 5GAN, in addition to all PDB codes listed in Extended Data Table 1 of [9]), mitochondrial ribosome (EMD 2490 and 2787; PDB 4CE4, 4V19, and 4V1A), CRISPR-Cas9-sgRNA complex (EMD 3276; PDB 5F9R, 4ZT0), U1 snRNP (EMD 8622; PDB 3CW1, 3PGW, 5GMK, 5UZ5), spliceosomal P complex (PDB 5MQ0, 5WSG, 5I8Q, 6BK8), HIV-1 RTIC (described in [44]), and *Tetrahymena* telomerase (EMD 6443; PDB 5KMZ, 2VOP, 5C9H, 2M21, 4ERD), MS2 packaged genome (EMD 8397 and 3403; PDB 5TC1). The DRRAFTER models of the U1 snRNP (from the 3.6 Å map) and the packaged MS2 genome (from the 3.6 Å map) are available in the Supplementary Information.

## Code availability statement

The DRRAFTER code is freely available to academic users as part of the Rosetta software package in weekly releases after March 14, 2018 at www.rosettacommons.org. Instructions for setting up Rosetta and running the DRRAFTER software are available at www.rosettacommons.org. A demo is also available at this site.

